# Efficient multi-task chemogenomics for drug specificity prediction

**DOI:** 10.1101/193391

**Authors:** Benoit Playe, Chloé-Agathe Azencott, Véronique Stoven

## Abstract

Adverse drug reactions, also called side effects, range from mild to fatal clinical events and significantly affect the quality of care. Among other causes, side effects occur when drugs bind to proteins other than their intended target. As experimentally testing drug specificity against the entire proteome is out of reach, we investigate the application of chemogenomics approaches. We formulate the study of drug specificity as a problem of predicting interactions between drugs and proteins at the proteome scale. We build several benchmark datasets, and propose *NN-MT*, a multi-task Support Vector Machine (SVM) algorithm that is trained on a limited number of data points, in order to solve the computational issues or proteome-wide SVM for chemogenomics. We compare *NN-MT* to different state-of-the-art methods, and show that its prediction performances are similar or better, at an efficient calculation cost. Compared to its competitors, the proposed method is particularly efficient to predict (protein, ligand) interactions in the difficult double-orphan case, i.e. when no interactions are previously known for the protein nor for the ligand. The *NN-MT* algorithm appears to be a good default method providing state-of-the-art or better performances, in a wide range of prediction scenarii that are considered in the present study: proteome-wide prediction, protein family prediction, test (protein, ligand) pairs dissimilar to pairs in the train set, and orphan cases.

## 1 Introduction

### 1.1 Drug specificity

The current paradigm in rationalized drug design is to identify a small molecular compound that binds to a protein involved in the development of a disease in order to alter disease progression. Once a hit ligand has been identified, often by combining *in silico* and *in vitro* approaches, this molecule needs to be optimized in order to meet the ADME (Absorption, Distribution, Metabolism, Elimination), toxicity, and industrial synthesis requirements. Finally, pre-clinical and clinical assays are organized to obtain agreement from the regulatory agencies. When successful, this process often lasts more than ten years, and recent estimates set the cost of drug development in US$2.5 billion in 2013 [1].

This complex, long, and costly process is often interrupted because of adverse drug reactions (ADR, also called side effects) that appear at various stages of drug development, or even after the drug has reached the market. In the USA, ADRs have been estimated to have an annual direct hospital cost of US$1.56 billion [2]. A meta-analysis estimated that, in hospitalized patients, the incidence of severe ADR was 1.9%- 2.3%, while the incidence of fatal ADR was 0.13% - 0.26%. For the year 1994, this means that 2 216 000 patients hospitalized in the US suffered from a serious ADR, and approximately 106 000 died [3]. Finally, a recent review [4] found that between 1950 and 2014, 462 medicinal products were withdrawn from the market in at least one country due to ADR. Of these 462 withdrawn drugs, 114 were associated with deaths.

Side effects frequently occur when drugs lack specificity, which means that they bind to proteins other than their intended target [5]. In that case, the molecular mechanisms at the source of the therapeutic effect and of the unwanted side effects are of similar nature: they both involve interactions between the drug and a protein. However, the complete study of drug specificity at early stages of drug development is experimentally out of reach, since it would require the evaluation of potential interactions between the hit molecule and the entire human proteome. Therefore, there is a strong incentive to develop *in silico* methods that predict specificity. The goal is to reduce the number of experiments to be performed, identify drug candidates that should be dropped because of their lack of specificity, protect patients from deleterious ADRs, and reduce the expense of time and money for the pharmaceutical industry.

### 1.2 Protein-ligand interactions prediction

The study of a drug’s specificity mainly boils down to predicting its protein targets in the space of the human proteome, or at least at the scale of “druggable” human proteins, i.e. proteins that present pockets into which drugs can bind. The approaches that have been developed to predict interactions between a protein and a small molecule can be separated into three categories.

First, *ligand-based approaches* such as Quantitative Structure Activity Relationship (QSAR) (refer to [6] for a recent review on QSAR) build a model function that predicts the affinity of any molecule for a given target, based on the affinities of known ligands for this target. They are efficient to study the affinity of molecules against a given protein target, but they are not suitable to study the specificity of a molecule against a large panel of proteins. This would indeed require, for each of the considered proteins, that the binding affinities of multiple ligands were available.

The second category is *docking* (refer to [7] for a recent review on docking), also called target-based approaches. Docking is a molecular modeling method that predicts the affinity of a ligand for a protein based on the estimated interaction energy between the two partners. However, it relies on the 3D structure of the proteins, which strongly limits its application on a large scale.

Finally, *chemogenomic approaches* [8] can be viewed as an attempt to fill a large binary interaction matrix where rows are molecules and columns are proteins, partially filled with the known protein-ligand interaction data available in public databases such as the PubChem database at NCBI [9]. In this context, drug specificity prediction is formulated as a classification problem, where the goal is to distinguish protein-ligand pairs that bind from those that do not: the aim is to predict “interacting” or “not interacting” labels for all pairs, but not to predict the strength of the interaction, which would correspond to a regression problem. Chemogenomics mainly belong to supervised machine learning (ML) methods, which learn mathematical models from available data, and use these models to make predictions on unknown data.

Various chemogenomics methods have been proposed in the last decade [10–24]. They all rely on the assumption that “similar” proteins are expected to bind “similar” ligands. They differ by (i) the descriptors used to encode proteins and ligands, (ii) how similarities between these objects are measured, (iii) the ML algorithm that is used to learn the model and make the predictions.

Predecessors of our approach include Support Vector Machine (SVM) [11], kernel Ridge Linear Regression (kernelRLS) [10, 12, 18–20], and matrix factorization (MF) [22–24].

MF approaches decompose the interaction matrix that lives in the (protein, molecule) space into the product of matrices of lower rank, living in the two latent spaces of proteins and of molecules. The most recent and efficient MF based approach by [24] consider more specifically Logistic Matrix Factorization [25]. They display good performances and are also computationally efficient. [24] also generalized their approach to orphan molecules and proteins by computing latent representations of orphan molecules and proteins as a weighted sum of the latent representation of their neighbors.

BLM make prediction for a (protein, molecule) pair based first on the prediction of target proteins for the considered molecule, and then on the prediction of ligand molecules for the considered protein. The predictor used is the kernelRLS. This gives two independent predictions for each putative drug-target interaction, which are combined into a final prediction.

Finally, kernel methods using the Kronecker product of the molecule and protein space (presented in the next section) can handle orphan cases, but are more computationally expensive. Among them, although the *KronRLS* method (a Kernel Regularized Least Square classifier) succeeded to dramatically reduce the computational complexity of its exact solution when used on the Kronecker product of the molecule and protein space, and is hence applicable to large scale chemogenomics studies, SVM-based methods are still computationally inappropriate at such scale. The present study propose a SVM-based approach and aims at addressing this issue.

In most cases, previous studies have been implemented to predict interactions of molecules with proteins belonging to the same family, such as kinases or GPCRs [10, 18–24, 26, 27]. A few studies have been devoted to larger scales in the protein space, such as [17] which however does not focus on settings relevant to the prediction of drug specificity. Some rely on the 3D structure of the binding pocket [28, 29], which limits the number of proteins that can be considered, others on coarse protein descriptors based on the presence of structural or functional domains [30]. In the present paper, we propose a computationally efficient approach to study the applicability of these ML techniques to the entire druggable proteome.

### 1.3 Single-Task and Multi-Task algorithms

In the context of the present paper, a *Single-Task* method consists in predicting protein targets for a given molecule *m*. In this setting, the specificity of *m* is studied by learning a model function *f*_*m*_(*p*) that predicts whether molecule *m* interacts with protein *p*, based on known protein targets for *m*. This means that a new model function is learned for each molecule. We refer to this setting as *ligand-based ST*. Conversely, a single-task method could learn a model function *f*_*p*_(*m*) that predicts whether protein *p* interacts with molecule *m*, based on all ligands known for protein *p*. We call this the *target-based ST* setting.

In contrast, *Multi-Task* methods predict whether *p* and *m* interact by training a model based on all known interactions, including those involving neither *m* nor *p*. In other words, the task of predicting ligands for protein *p* is solved not only based on the data available for this task (i.e. known ligands for this protein), but also based on the data available for the other tasks (i.e. known ligands any other protein). The main issue is to define how the data available in all tasks can be used to make the predictions made for a given task.

More generally, the main idea behind multi-task learning is that, when solving several related tasks for which the data available for each task are scarce, a multi-task framework defines how to share information across tasks, which can improve the performance of the final prediction models that can be built.

The concept of multi-task learning is related to that of transfer learning [85]. Transfer learning is inspired from the fact that humans are able to use their acquired knowledge and experience to solve new problems faster. More precisely, according to the classification by [31], our multi-task learning problem falls into the category of inductive transfer learning, meaning that the training and testing domains are the same, i.e. the training and testing data are encoded in the same space, and the tasks are different but related. In our case, we predict protein-ligand interactions by training a model based on known protein-ligand interactions (i.e. same space for training and tested data), and the tasks are predictions of ligands for proteins whose sequences can be compared (i.e. the tasks are related). Such approaches are of particular interest when the data available for each task are scarce, i.e. when few interactions are known for a given ligand or protein, as is often the case when looking for secondary targets at the size of the human proteome. Even within the multi-task framework, orphan proteins (for which no ligand is known) and proteins for which the only known ligands are very dissimilar to the tested molecule, are those for which predictions are the most difficult. Therefore, our study will focus more particularly on these cases.

Among all multi-task approaches, multi-task kernel methods have widely been used in bioinformatics, including for chemogenomics applications. Our contributions in the present paper belong to this category of methods that we briefly review in the next section.

### 1.4 Kernel methods for chemogenomics

In this study, we formalize and solve the problem of drug-target interaction prediction with Support Vector Machines (SVM), an algorithm for learning a classification or a regression rule from labeled examples [32].

Intuitively, SVMs seeks to find the optimal hyperplane separating two classes of data points. As briefly recalled in Supplementary Materials S1, although SVMs can be solved from vector representations of the data, they can also be solved using the “kernel trick”, based only on the definition of a kernel function *K* which gives the similarity value *K*(*x, x*^′^) between all pairs of data points *x* and *x*^′^, without needing an explicit representation of the data. Many kernels have been proposed for molecules and for proteins, and an overview of such kernels is presented in the Material and Methods section.

In chemogenomics, our goal can be viewed as finding the optimal hyperplane that separates the pairs (*m, p*) of molecules and proteins that interact from those that do not interact. This classification task can be solved using an SVM with a kernel *K*_*pair*_ defined on (ligand, protein) pairs. Given *N* example pairs, solving the SVM in the space of (*m, p*) pairs using the *K*_*pair*_ kernel corresponds to finding the optimal *α*_*i*_ coefficients such that (see Supplementary Materials S1):

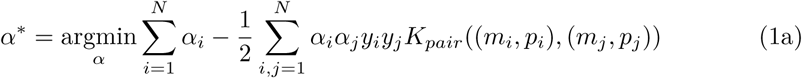

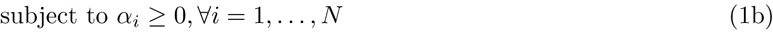

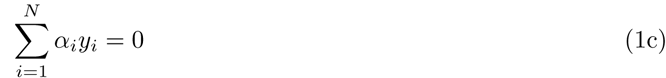

A general method to build a kernel on such pairs is to use the Kronecker product of the molecule and protein kernels [33]. Given a molecule kernel *K*_*molecule*_ and a protein kernel *K*_*protein*_, the Kronecker kernel *K*_*pair*_ is defined by:

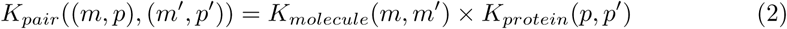

Thus, the Kronecker kernel *K*_*pair*_ captures interactions between features of the molecule and features of the protein that govern their interactions (see Supplementary Materials S2 for an explicit definition of the Kronecker product of two matrices). If *K*_*molecule*_ is a *n × n* matrix and *K*_*protein*_ is a *p × p* matrix, their Kronecker product *K*_*pair*_ has size *np × np*. In the context of chemogenomics, this can correspond to a very large size, leading to untractable computations. However, one interesting property of the Kronecker kernel is that calculating its values on a data set of (*m, p*) pairs does not require storing this entire matrix since it is sufficient to store *K*_*molecule*_(*m, m*^′^) and *Kprotein*(*p, p*^′^).

Therefore, solving the SVM (equation 1) only requires calculation of the *K*_*molecule*_ and *K*_*protein*_ kernels according to equation 2.

Once the *α*_*i*_ coefficients have been determined, the ability of a given (*m, p*) pair to interact is predicted based on:

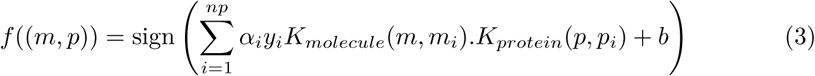

This equation illustrates why the use of such a kernel can be viewed as a multi-task method. Indeed, in a single-task approach where one task corresponds to the prediction of ligands for a given protein *p*, the ability of molecule *m* to bind protein *p* would be estimated by:

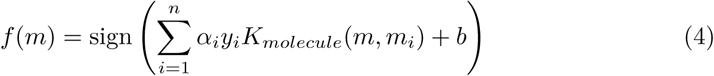

where the *m*_*i*_ molecules are ligand and non-ligand molecules known for protein *p*.

In the multi-task setting, *f* ((*m, p*)) evaluates the ability of *m* to bind to protein *p* using *m*_*i*_ molecules that are all ligand or non-ligand molecules known for all proteins *p*_*i*_. However, the contribution of the labels *y*_*i*_ of ligands for *p*_*i*_ proteins that are different from *p* to calculate *f* ((*m, p*)) is weighted by *K*_*protein*_(*p, p*_*i*_). In other words, the more similar two tasks (i.e. the corresponding proteins) are, the more known instances for one of the task will be taken into account to make predictions for the other task.

Such kernel-based multi-task approaches have been successfully applied to biological problems, including the prediction of protein-ligand interactions [8, 11, 34, 35].

### 1.5 Contribution

The goal of this paper is to investigate the application of multi-task Support Vector Machines (SVM) to the prediction of drug specificity, by predicting interactions between the drug of interest and the entire druggable proteome. To this end, we evaluate multi-task SVMs in several key scenarii that explore the impact of the similarity between the query (protein, molecule) pair and the training data on the prediction performance, a point that is rarely discussed in the literature. We also explore their applicability to orphan settings, a situation often encountered large scale studies, and where single-task methods are not applicable. We also discuss how to generate negative training examples, and we optimize protein and molecule kernels in the spaces of drug-like molecules and druggable proteins.

Our observations lead us to propose the *NN-MT* algorithm, a multi-task SVM for chemogenomics that is trained on a limited number of data points: for a query (protein, molecule) pair (*p*^*\**^, *m*^*\**^), the training data is composed of (1) all *intra-task* (protein, ligand) pairs defined by pairs (*p, m*) with either *p* = *p*^*\**^ or *m* = *m*^*\**^; (2) a limited number of *extra-task* (protein, ligand) pairs, defined by pairs (*p, m*) with *p* ≠ *p*^*\**^ and *m* ≠ *m*^*\**^, chosen based on the similarity of *p* and *m* to *p*^*\**^ and *m*^*\**^, respectively; and (3) randomly picked negative examples (about ten times more than positive training pairs). While the applicability of multi-task approaches can be limited in practice by computational times, our approach only requires training on a dataset of size similar to those used by single-task methods. We evaluate the performance on various assembled datasets in which the protein and/or the ligand are orphan.

We also compare the *NN-MT* algorithm to state-of-the-art approaches in drug-target interaction prediction [18–24]. We used and updated the PyDTI package [24], adding an implementation of *NN-MT* together with key cross-validation schemes and a DrugBank-based dataset built in the present study. In addition to all other experiments performed in the present study, this benchmark study concludes that *NN-MT* is a good default method providing state-of-the-art or better performances, in a wide range of prediction scenarii that can be encountered in real-life studies: proteome-wide prediction, protein family prediction, test (protein, ligand) pairs dissimilar to pairs in the train set, and orphan cases.

All datasets and codes are available at https://github.com/bplaye/efficient_MultiTask_SVM_for_chemogenomics/. The updated PyDTI package is available at https://github.com/bplaye/PyDTI/.

## 2 Materials and methods

### 2.1 Protein kernels

We used sequence-based kernels since they are suitable for proteome-wide approaches, unlike kernels relying on the 3D structure of the proteins or on binding pocket descriptions. Numerous studies have already been devoted defining descriptors of proteins based on amino-acid sequence [36–40]. We considered three sequence-based kernels: the *Profile kernel* [40], *the SWkernel*, and the *LAkernel*.

The *Profile kernel* uses as protein descriptors the set of all possible subsequences of amino acids of a fixed length *k*, and considers their position-dependent mutation probability. This kernel is available at http://cbio.mskcc.org/leslielab/software/string_kernels.html.

We also used two kernels that rely on local alignment scores. The first one is directly based on the Smith-Waterman (SW) alignment score between two proteins [41] and is called the *SWkernel* in the present paper. SW scores were calculated with the EMBOSS Water tool available at http://www.ebi.ac.uk/Tools/psa/emboss_water/. We built a kernel based on the SW score matrix by subtracting its most negative eigenvalue from all diagonal values. We also used the Local Alignment kernel (*LAkernel*) [39] which mimics the behavior of the SW score. However, while the SW score only keeps the contribution of the best local alignment between two sequences to quantify their similarity, the LAkernel sums up the contributions of all possible local alignments, which proved to be efficient for detecting remote homology [39]. This kernel is available at http://members.cbio.mines-paristech.fr/jvert/software/.

#### Kernel hyperparameters values

The Profile kernel has two hyperparameters: the size *k* of the amino acid subsequences that are searched and compared, and the threshold *t* used to define the probabilistic mutation neighborhoods. We considered *k* ∈ {4, 5, 6, 7} and *t* ∈ {6, 7.5, 9, 10.5}. The SWkernel also has two hyperparameters: the penalties for opening a gap (*o*) and for extending a gap (*e*). We considered *O* ∈ {1, 10, 50, 100} and *e* ∈ {0.01, 0.1, 0.5, 1, 10}. The LAkernel has three hyperparameters: the penalties for opening (*o*) and extending (*e*) a gap, and the *β* parameter which controls the importance of the contribution of non-optimal local alignments in the final score. We considered *o* ∈ {1, 20, 50, 100}, *e* ∈ {0.01, 0.1, 1, 10}, and *β* ∈ {0.01, 0.5, 0.05, 0.1, 1}. All kernels hyperparameters were optimized by cross-validation (see Section 2.3).

In the last part of the study, we also considered kernels on proteins based on their family hierarchy. Indeed, the most important classes of drug targets have been organized into hierarchies established on the sequence and the function of the proteins within these families (GPCR [42], kinases [43] and ion channels [44]). As in [11], the hierarchy kernel is built based on the number of common ancestors shared by two proteins in the hierarchy. More precisely, *K*_*hierarchy*_(*t, t*^′^) = 〈 *ϕ*(*t*), *ϕ*(*t*^′^) 〉, where *ϕ*(*t*) is a binary vector for which each entry corresponds to a node in the hierarchy and is set to 1 if the corresponding node is part of *t*’s hierarchy and 0 otherwise.

All protein kernels were centered and normalized.

### 2.2 Small molecule kernels

Many descriptors have been proposed for molecules, based on physico-chemical and structural properties [45–48]. To measure the similarity between molecules, we considered two state-of-the-art kernels based on molecular graphs that represent the 2D structure of the molecules, with atoms as vertices and covalent bonds as edges. Both kernels compute similarities between molecules via the comparison of linear fragments found in their molecular graphs. They are available at http://chemcpp.sourceforge.net/.

The first one, called the Marginalized kernel [47], calculates the similarity between two molecules based on the infinite sets of random walks over their molecular graphs.

The second kernel, called the Tanimoto kernel, uses a description of molecules by vectors whose elements count the number of fragments of a given length. The similarity between molecules is based on the Tanimoto metric [45].

#### Kernel hyperparameters values

The Marginalized kernel has two hyperparameters: the stopping probability (while building a path) *q* in {0.01, 0.05, 0.1, 0.5}, and the Morgan Index (MI) in {2, 3, 4}. For both kernels, hyperparameters were selected by cross-validation (see Section 2.3). The Tanimoto kernel has one hyperparameter: the length *d* of the paths, which we considered in {2, 4, 6, 8, 10, 12, 14}. All molecule kernels were centered and normalized.

### 2.3 Evaluation of prediction performance

Prediction performance is commonly evaluated with a cross-validation (CV) scheme [49]: 1) the dataset is randomly split into K folds 2) the model is run K times, each run using the union of (K-1) folds as the training set, and measuring the performance on the remaining fold. Prediction performance are averaged over all folds. When hyperparameters had to be selected, we used a nested cross validation (*Nested-CV*) scheme [50]. It consists in a (K-1) folds cross validation (*inner-CV*) nested in a K folds cross validation (*outer-CV*). At each step of the outer-CV, the inner-CV is repeated for all considered values of the hyperparameters. The values leading to the best prediction performance are retained as optimal. We used K=5, a classical value in CV.

We also considered leave-one-out cross-validation (*LOO-CV*), for which the number of folds is the number of available points in the dataset. The *LOO-CV* scheme is particularly useful when the number of samples is small. It was used in the present paper when the size of the considered dataset was too small to perform *5-fold-CV*.

We estimated prediction performance using two scores that are classically employed to judge the quality of a classifier in case of drug-target interaction prediction. The first one is the area under the Receiver Operating Characteristic curve [51] (ROC-AUC). The ROC curve plots true positive rate as a function of false positive rate, for all possible thresholds on the prediction score. Intuitively, the ROC-AUC score of a classifier represents the probability that if a positive and a negative interaction are each picked at random from the dataset, the positive one will have a higher positive score than the negative one. The second one is the area under the Precision-Recall curve [52] (AUPR). It indicates how far the prediction scores of true positive interactions are from false positive interactions, on average. Although we used both the ROC-AUC and AUPR scores, since negative interactions are actually unknown interactions in protein-ligand interaction datasets, the AUPR is considered a more significant quality measure of the prediction method than the ROC-AUC. Indeed, it emphasizes the recovery of the positive samples and penalizes the presence of false positive examples among the best ranked points.

We used the Python library scikit-learn [53] to implement all considered machine learning algorithms.

### 2.4 Datasets

Many publicly available databases such as KEGG Drug [54], DrugBank [55], or ChEMBL [56] can be used to build a learning dataset of protein-ligand interactions. We chose to build all the datasets used in the present study from the DrugBank database v4.3, because it contains FDA-approved drugs, or drug candidate molecules. This allowed optimize and test our models on drug-like molecules, on which they intend be applied. In addition, we assumed that the list of human proteins appearing as targets for molecules of DrugBank can represent a relevant “druggable” human proteome on which we could train models that predicting the specificity of drug-like molecules.

We built a first learning dataset called *S*, based on Version 4.5 of the DrugBank [55]. We selected all molecules targeting at least one human protein, and having a molecular weight between 100 and 600 g.mol^-1^, a range in which most small molecule marketed drugs are found [57]. This leads to a dataset composed of 3 980 molecules targeting 1 821 proteins, and including 9 536 protein-ligand interactions that correspond to the positive training pairs. All other protein-ligand pairs are unlabeled because no interactions were recorded for them in the database. Most of these pairs are expected not to interact, but a small number of them are in fact missing interactions. However, we considered that all unlabeled pairs as negative examples, allowing the predictor to re-classify some of these pairs as positive examples.

We built several other datasets using exactly the same training pairs as those in *S*, but 5-folded in various ways. Datasets *S*_1_, *S*_2_, *S*_3_, and *S*_4_ are folded so as to correspond to random, orphan protein, orphan ligand, and double orphan prediction situations. The construction of these four datasets is detailed in Section 3.2, where they are used. Datasets 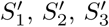 and 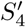 are also folded to mimic the same situations, but with the additional constraint that proteins and ligands were clustered based on their similarities, and each fold contains only one cluster of proteins and of ligands. The goal is to test the performance of the method in situations similar to those of *S*_1_, *S*_2_, *S*_3_, and *S*_4_, but with the added difficulty that the test set (one fold) and the train set (the 4 other folds) contain pairs that have low similarities. This setting is relevant when considering proteome-wide predictions: many of the proteins to consider may not have close neighbors among the proteins for which the most information (i.e. ligands) are known. The construction of these four datasets is detailed in Section 3.3, where they are used.

We also built a dataset called *S*_0_ by keeping only molecules and proteins in *S* which are involved at least in two interactions, in order to compare the prediction performance of the proposed methods with those of ligand-based and target-based approaches. Indeed, these two single-task approaches require at least two data points, one used as train, and one as test. Consequently, when a *LOO-CV* scheme is used, no ligand and no protein are orphans in *S*_0_. *S*_0_ contains 5 908 positive interactions and was used in Sections 3.4 and 3.5. In addition, we randomly generated four sets of 5 908 negative interactions involving proteins and ligands found in the positive interactions, while ensuring that each protein and each ligand are present in the same number of positive and negative interactions. Then, we assessed performance by computing the mean and standard deviation of the AUPR scores over test sets including the positive interactions set and one of the negative interactions sets.

Finally, we built three protein family datasets by extracting from *S*_0_ all protein-ligand interactions involving respectively only G-Protein Coupled Receptors (GPCR set), ion channels (IC set), and kinases (Kinases set). These datasets were used to evaluate performance of our method within a family of proteins, and compare it to those of single-task approaches. They were extracted from *S*_0_ (and not from the larger dataset *S*) since again, these comparisons used the *LOO-CV* scheme, which requires at least two data points per protein and per molecule.

Table 1 gives some statistics about the datasets, including the distribution of targets per drug and the distribution of ligands per protein.

**Table 1.** Dataset statistics

| | $S$ | $S_0$ | GPCR | IC | Kinases |
| --- | --- | --- | --- | --- | --- |
| number of interactions | 9 536 | 5 908 | 1 735 | 1 603 | 847 |
| number of proteins | 1 821 | 788 | 85 | 140 | 143 |
| number of molecules | 3 980 | 1 180 | 482 | 295 | 577 |
| number of targets per drug (mean/median) | 5.2/2 | 7.5/3 | 20.3/6 | 5.9/3 | 11.5/4 |
| number of targets per drug (min – max) | 1 – 136 | 2 – 82 | 1 – 86 | 1 – 67 | 1 – 136 |
| number of ligands per protein (mean/median) | 2.4/1 | 5.0/3 | 3.6/3 | 5.4/3 | 1.5/7 |
| number of ligands per protein (min – max) | 1 – 70 | 2 – 48 | 1 – 31 | 1 – 26 | 1 – 18 |

## 3 Results

### 3.1 Kernel selection and parametrization

The goal of this section is to choose a protein kernel and a molecule kernel that we will use throughout the remainder of this study. We assumed that kernels optimized on a large dataset of interactions between drug-like molecules and druggable human proteins such as dataset *S* would be good default kernels for the prediction of drug candidates specificity. Therefore, we optimized kernels on dataset *S* (the largest dataset built in the present study), and used the best-performing couple of kernels in the remainder of the paper.

The set of (protein, ligand) pairs in *S* were randomly 5-folded, and we performed a *nested 5-fold-CV* experiment in order to evaluate the six possible kernel combinations and their best hyperparameters.

Table 2 gives the best prediction performance for the six combinations of protein and molecule kernels, together with the corresponding hyperparameters. All protein kernels gave the best AUPR when coupled to the Tanimoto kernel. The Marginalized kernel obtained good performance only when coupled to the Profile kernel. Overall, the Profile kernel (*k* = 5, *t* = 7.5) associated to the Tanimoto kernel (*d* = 8) gave the best performance. Therefore, in what follows, we only consider these two kernels, and call *MT* the Multi-Task SVM that uses their Kronecker product.

**Table 2.**
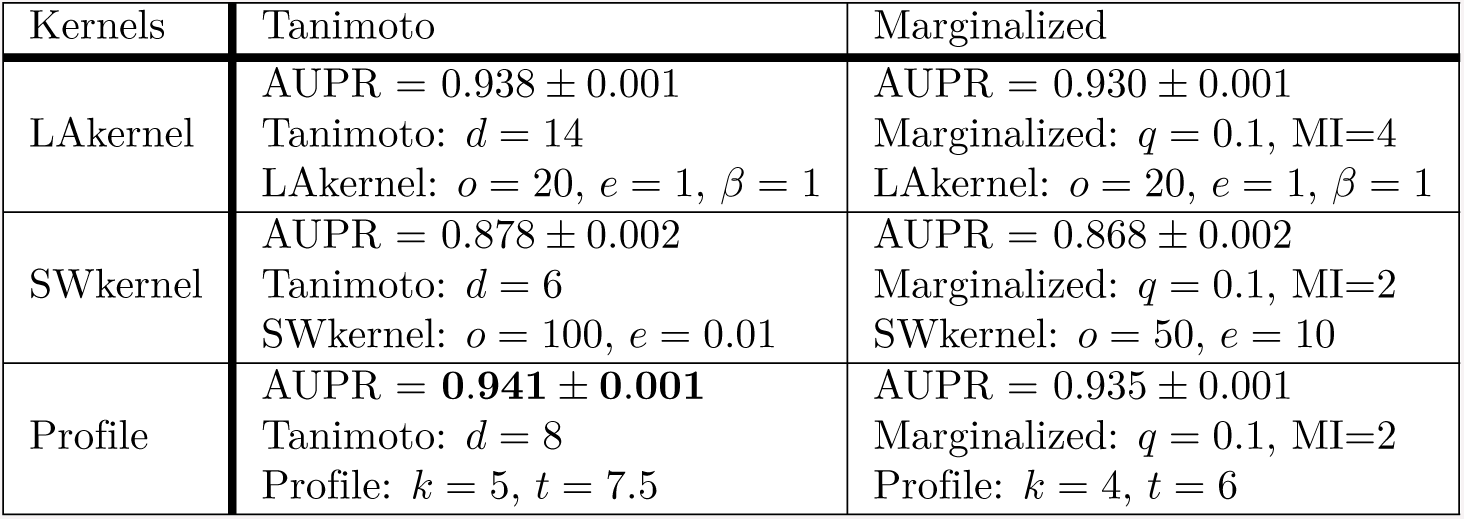
Best *nested 5-fold-CV* AUPR for each kernel combination, together with optimal hyperparameters.

We also considered one-class SVM using the same kernels [58]. However, the performance of one-class SVM were clearly lower than those of KronSVM. The AUPR scores of one-class SVM were in the range of 0.6 for all considered kernels when those of KronSVM were in the range of 0.9. Therefore, we did not further consider one-class SVM.

It is worth noting that the SW-kernel gave the worst performance, although it is used in many studies [10, 15, 18, 20]. Overall, the good performance of the six multi-task methods observed on *S* is consistent with previously reported results [17, 20]. However, *S* was built from the DrugBank, which is mostly fueled by application-specific screens of either related proteins or related small molecules. Therefore, (protein, ligand) pairs of the test sets will usually have close pairs in the train set (i.e. pairs involving the same or similar proteins and ligands), which will facilitate the prediction. The performance in real-case prediction of drug specificity is expected to be lower than that obtained on *S*, since at the proteome scale, some of the test (protein, ligand) pairs will be far from all pairs of the train set. This will be particularly true in the case of new drugs and therapeutic targets, as already pointed by [26].

The question of interest is now to which extent the proposed *MT* method is effective to make predictions on more challenging situations that are relevant in the context of drug specificity prediction. Therefore, in the following, we study the evolution of *MT*’s performance in more realistic settings where the protein, the molecule, or both, are orphan, or where the tested (protein, ligand) pair has low similarity with the pairs belonging to the train set.

### 3.2 Performance of multi-task approaches in orphan situations

The goal of this section is to evaluate the performance of *MT* in cases where the queried (protein, molecule) pairs contain proteins and/or molecules that are *not* in the training set, as proposed by [26]. For that purpose, all the pairs of dataset *S* were used and folded as follows in order to create four cross-validation data sets:

- *S*_1_: randomly and balanced in positive and negative pairs;
- *S*_2_ (corresponding to the “orphan ligand” case): (protein, molecule) pairs in one fold only contain molecules that are absent from all other folds; prediction on each test set (each fold) is performed using train sets (the four other folds) in which no the ligands of the test set are absent.
- *S*_3_ (corresponding to the“orphan protein” case): (protein, molecule) pairs in one fold only contain proteins that are absent from all other folds; prediction on each test set is performed using train sets in which no the proteins of the test set are absent.
- *S*_4_ (corresponding to the“double orphan” case): (protein, molecule) pairs in one fold only contain proteins *and* molecules that are both absent from all other folds. Prediction on each test set is performed using train sets in which no the proteins and the ligands of the test set are absent. The folds of *S*_4_ were built by intersecting those of *S*_2_ and *S*_3_ and *S*_4_. Thus, *S*_4_ contains 25 folds and not 5.

Fig 1 shows the nested-CV AUC and AUPR scores obtained by the *MT* method on the *S*_1_-*S*_4_ datasets. As expected, the best scores are obtained for *S*_1_, and the worst for *S*_4_, since in *S*_4_, no pairs of the train set contain the protein or the ligand of the tested pair to guide the predictions. The loss of performance between the random and the double orphan settings is about 0.12 both in AUC and AUPR. However, the performance on the *S*_4_ dataset remains well above those of a random predictor. These results confirm that *MT* chemogenomics can make predictions for (protein, ligands) pairs made of unseen proteins and unseen ligands, even in datasets containing very diverse types of proteins. This confirms previous observations made on less diverse datasets [26]. It is important to point that single-task approaches would not be able to provide any prediction on the *S*_4_ dataset.

**Fig 1.** *nested 5-fold-CV* **performance of the** *MT* **method on the** *S*_1_ *-S*_4_ **datasets.** Numerical values can be found in Supporting Information S1 Table.

The scores obtained in the *S*_2_ and *S*_3_ datasets are intermediate between those observed on *S*_1_ and *S*_4_. This was to be expected, as in these datasets, the algorithm can rely on training pairs containing either the same proteins (*S*_2_) or the same ligands (*S*_3_) as the test set. The AUC and AUPR scores are both slightly better for *S*_3_ than for *S*_2_, which suggests that predicting ligand for new protein targets is easier than predicting targets for new compounds, as already noticed in [26]. We also observed similar behaviors when replacing the SVM with a kernel ridge regression (see Supporting Information S1 Fig) and hence did not further consider this algorithm.

Overall, our results suggest that the performance of *MT* is driven by known (protein, molecule) pairs that are similar to the query pair, in the sense that they share either their protein or their molecule. In the next section, we will evaluate how the actual similarity between query and train pairs influences the prediction performance of this multi-task algorithm.

### 3.3 Impact of the similarity of the training examples to the test set

To evaluate the impact on performance of the dissimilarity between training and test pairs, we re-folded the pairs of *S* following the “clustered cross-validation” approach [59]. More precisely, we clustered proteins (resp. ligands) into 5 clusters by hierarchical clustering [60]. We then built four cross-validation datasets, 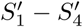, generated based on folds similarly as *S*_1_ *-S*_4_, but with the added constraint that all pairs in a given fold are made of proteins from a single protein cluster and ligands from a single ligand cluster. Therefore, test pairs are more dissimilar from train pairs than in the *S*_1_ *-S*_4_ datasets, which makes the problem more difficult.

Overall, all the pairs of dataset *S* were 5-folded as follows in order to create four cross-validation data sets:

- 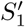: randomly and balanced in positive and negative pairs, each fold containing proteins and ligands belonging to the same cluster;
- *S*_2_’ (corresponding to the “orphan ligand” case): (protein, molecule) pairs in one fold only contain molecules that are absent from all other folds; prediction on each test set (each fold) is performed using train sets (the four other folds) in which no the ligands of the test set are absent, with the additional constraint that each fold contains proteins and ligands belonging to the same cluster.
- 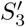 (corresponding to the “orphan protein” case): (protein, molecule) pairs in one fold only contain proteins that are absent from all other folds; prediction on each test set is performed using train sets in which no the proteins of the test set are absent, with the additional constraint that each fold contains proteins and ligands belonging to the same cluster.
- 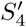 (corresponding to the “double orphan” case): (protein, molecule) pairs in one fold only contain proteins *and* molecules that are both absent from all other folds. Prediction on each test set is performed using train sets in which no the proteins and the ligands of the test set are absent, with the additional constraint that each fold contains proteins and ligands belonging to the same cluster. The folds of *S*_4_ were built by intersecting those of *S*_2_ and *S*_3_ and *S*_4_. Thus, *S*_4_ contains 25 folds and not 5.

**Fig 2.** *nested 5-fold-CV* **performance of the** *MT* **method on the 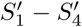 datasets.** Numerical values can be found in Supporting Information S2 Table.

We used the same kernels as for the *MT* method. Fig 2 shows the prediction performance of *MT* on these new cross-validation folds. For all the datasets, we observed a strong decrease in prediction scores with respect to those obtained on the corresponding *S*_1_ *-S*_4_ datasets. This suggests that good performance on a query pair (*p*^*\**^, *m*^*\**^) is driven by the presence in the training set of pairs made *both* of proteins similar to *p*^*\**^ and of molecules similar to *m*^*\**^, even if the query pair (*p*^*\**^, *m*^*\**^) is a double orphan, as in *S*_4_. Our results also suggest that it is more important to train on pairs similar to the double orphan query pair (*p*^*\**^, *m*^*\**^), as in *S*_4_, than on data containing, for example, *p*^*\**^ itself, but paired only with molecules quite dissimilar to *m*^*\**^, as in 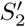.

Finally, performance on 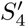 are random, confirming that making predictions for double orphans that are also very dissimilar from the training data is a very difficult task.

These results suggest that pairs in the training set that are very dissimilar to the query pair do not help making more accurate predictions. In other words, although the kernels used in multi-task approaches modulates how information available in one task is shared for training other tasks (the further the tasks are, the less information is shared), using information from distant tasks seems to degrade performance. This insight is interesting since the *MT* algorithm requires calculating the Kronecker kernel on all (protein, molecule) pairs, which is computationally demanding. Therefore, the next two sections evaluate whether removing distant pairs from the training set can improve computational efficiency without degrading performance.

### 3.4 Multi-task approaches on reduced training sets

Based on the insight that *MT* prediction is driven by training examples that are close to the query data, we propose to build training tests of reduced sizes by removing training examples distant from the query. The goal of this section is also to compare the prediction performance of the *MT* method trained on these reduced data sets to that of the simpler and faster single-task method, since there would be no point in using the more complex *MT* method if a single-task method performs better. Because this study is motivated by ligand specificity prediction, we chose to focus on comparisons with the *ligand-based ST* method rather than *target-based ST*.

In what follows, *n*^+^ (resp. *n*^-^) will refer to the number of positive (resp. negative) examples in the train set.

In all the following experiments, we used the *LOO-CV* scheme because intra-task and extra-task pairs can only be defined for each pair separately, which prevents from using *K-fold-CV* schemes. In addition, in single-task approaches, the size of the training set was relatively small in most cases (see datasets statistics in Section 2), which does not allow to fold the data. We checked that the *LOO-CV* scheme did not trigger a bias, as sometimes observed [26] (see Supporting Information S2 Fig and S3 Table).

Because prediction of a given (protein, ligand) interaction can only be made by single-task if the pair partners are present in at least another pair of the train set, in the following experiments, we used the *S*_0_ dataset in which all ligands and all proteins are involved in at least two known interactions, as explained in Section 2.4.

#### 3.4.1 Training on intra-task positive examples

The goal of this section is to compare the prediction performance of the *MT* method trained on a reduced data set (of size similar to that employed in single-task methods) to those of single-task methods. Since *ligand-based ST* can only use intra-task positive examples, the only positive training pairs we use for the *MT* method are the intra-task pairs as well. Note that *MT* still gets more training examples than *ligand-based ST*, since pairs formed with the query protein and a different ligand are also included. By reducing the training set size, the computational times required by the *MT* method are now similar to those of the single-task method. In the following, we call *MT-intra* this variant of *MT*. For each test ligand, we build the negative training examples by randomly selecting a number *n*^-^ of proteins that do not interact with the ligand in *S*_0_. We vary *n*^-^ from 1 to 100 *× n*^+^.

Fig 3 shows the *LOO-CV* AUPR of *MT-intra* and *ligand-based ST* on *S*_0_, for increasing values of the *n*^-^*/n*^+^ ratio. For both methods, the AUPR score increases with the number of negative pairs in the train set, before decreasing for large numbers of negative pairs. A good trade-off for both computational and predictive performance seems to be in the range of 10 times more negative points than positive points. We therefore set *n*^-^*/n*^+^ to 10 for the remaining experiments of this section.

The AUPR scores of *MT-intra* outperform those of the *ligand-based ST* method. Interestingly, the performance of the *MT-intra* with a *n*^-^*/n*^+^ ratio of 10 is close to 0.96 which outperforms the AUPR score of 0.93 obtained with *MT* (see Section 3.2). This indicates that including in the train set pairs displaying low similarity to the tested pair degrades both the computational time and the quality of the prediction of *MT*.

**Fig 3.** Performance of single-task and *MT-intra* **as a function of the** *n*^-^*/n*^+^ **ratio**. Numerical values can be found in Supporting Information S4 Table

#### 3.4.2 Adding similar extra-task positive examples

The results from Section 3.3 suggest to explore the performance of the *MT-intra* method when trained on various datasets including extra-task pairs close to the tested pair, in addition to the intra-task pairs. Therefore, we built train sets made of:

- the train set of *MT-intra*
- 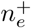 closest extra-task positive pairs with respect to the tested pair 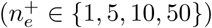.
- 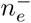 closest extra-task negative pairs with respect to the tested pair, so that the 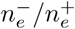 ratio varies from 1 to 10.

We call *NN-MT* (for Nearest Neighbor *MT*) the resulting variant of *MT*. We also considered a similar approach in which the extra-task pairs were chosen at random rather than according to their similarity to the test pair. We refer to this method as *RN-MT* (for Random Neighbor *MT*).

We report the *LOO-CV* performance of *NN-MT* and *RN-MT* on Fig 4. Fig 4(a) shows that, while adding to the train set 0 to 50 nearest neighbor extra-task positive pairs with respect to the tested pair, the prediction performance of *NN-MT* slightly and monotonously increases. Fig 4(b) shows that the performance of *RN-MT* also slightly increases (although not monotonously) when random extra-task pairs are added. However, its best performance remains under that of *NN-MT*. Finally, using a high 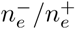 ratio did not improve the performance. This is an interesting observation, since limiting the size of the train set is computationally favorable.

**Fig 4.** AUPR as a function of the 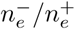 ratio for increasing numbers random extra-task points in the train set. (a): *NN-MT*. (b): *RN-MT*. The blue horizontal line corresponds to *MT-intra* (which is trained only on intra-task pairs). Numerical values can be found in Supporting Information S5 and S6 Tables

Taken together, these results show that *NN-MT* outperforms not only *MT-intra*, but also the more computationally demanding *MT* method trained in the *LOO-CV* setting in Section 3.2. AUPR scores for (protein, ligand) pairs involving non-orphan ligands and non-orphan proteins are expected to be very high (around 0.96).

However, predicting the specificity of a given ligand requires the ability to make predictions for proteins that are far from the known targets. In these cases, the high prediction scores obtained in this section might not hold. Therefore, in the next section, we study the performance of *NN-MT* when the test pairs are far from the train set.

### 3.5 Impact of the distance of the intra-task examples to the query pair

The goal of this section is to evaluate the performance of the *MT-intra* and *NN-MT* proposed methods, and to compare them to those of the *ligand-based ST* method, when the similarity between the test pair and the training data varies.

#### 3.5.1 Training on dissimilar intra-task positive examples

We first evaluated the performance of *ligand-based ST* and *MT-intra* when the similarity between the test pair and the training data varies. To do so, we computed the percentiles of the molecules (respectively proteins) similarity distribution in *S*_0_.

For each test pair (*p*^*\**^, *m*^*\**^), the training set only included the positive intra-task pairs (*p, m*) such that *K*_*protein*_(*p, p*^*\**^) and *K*_*molecule*_(*m, m*^*\**^) is lower than a percentile-based threshold *θ*. We then added *n*^-^ random intra-task negative pairs. We did not apply a similarity constraint to negative pairs, since, unlike the positive pairs, they are available in large numbers and at all distances from the tested pairs.

Fig 5 reports the *LOO-CV* AUPR scores of *ligand-based ST* and *MT-intra* for varying values of *θ* (20^*th*^, 30^*th*^, 50^*th*^, and 80^*th*^ percentiles) and of the *n*^-^*/n*^+^ ratio (from 1 to 50).

Fig 5(a) shows that, as expected, the performance of *MT-intra* increases when the similarity of the tested pair to the train set increases from the 20th to the 80th percentiles (AUPR of 0.67 to 0.77). However, the performance is still much lower than when the closest pairs are allowed in the training set (AUPR of 0.96, see Section 3.4.1). Fig 5(a) also suggests that a *n*^-^*/n*^+^ ratio of 10 is again an appropriate choice, as when all intra-task positive example are used (see Section 3.4.1). We therefore set *n*^-^*/n*^+^ to this value for the remaining experiments of this section.

**Fig 5.** AUPR scores as a function of the *n*^-^*/n*^+^ **ratio, for percentile-based threshold** *θ* **ranging from 20% to 80%.** (a): *MT-intra* method. (b): *ligand-based ST* method. Numerical values can be found in Supporting Information, respectively S7 and S8 Tables

Fig 5(b) shows that *ligand-based ST* behaves similarly to *MT-intra*: the AUPR score increases from 0.70 to 0.75 for *ligand-based ST* when threshold *θ* increases from the 20th to the 80th percentiles. These values again remain much under the AUPR score of 0.93 observed when all intra-task pairs are used. Although modest, the performance of *MT-intra* remains above those of *ligand-based ST* for all tested thresholds of similarity (except *θ* = 20) between the tested pair and pairs of the train set.

### 3.5.2 Adding similar extra-task positive examples

We then explored to which extent adding extra-task (protein, ligand) pairs to the training set of the *MT-intra* method improves the prediction scores.

Applying the same percentile-based similarity constraint to the intra-task positive pairs, we compared the performance of *NN-MT* and *RN-MT* when respectively adding 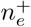 nearest neighbors or random extra-task positive to the training set. We did not apply a similarity constraint to the extra-task pairs, since the principle underlying multi-task methods is precisely to learn from extra-task data, which is particularly critical when the intra-task pairs of the train set are scarce or far from the tested pair, as illustrated by the poor performance of *ligand-based ST* in the previous section. A number 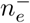 of nearest neighbors (respectively random) negative extra-task pairs were also added for *NN-MT* (respectively *RN-MT*).

Figs 6(a) and (b) report the *LOO-CV* AUPR of *NN-MT*, as a function of 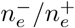 (ratio of negative over positive extra-task pairs) and for a number of extra-task positive pairs varying from 0 to 50, respectively for percentile similarity constraints *θ* of 20 and 80. The blue horizontal line (for 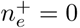) corresponds to the performance of the *MT-intra* methods. Fig 6(a) and (b) show that adding extra-task pairs to the train set dramatically improves performance. The AUPR score reaches values above 0.85, independently of *θ*, suggesting that when no close intra-task pairs are available, performance is driven mainly by extra-task training pairs, confirming our observations in Section 3.3. Moreover, when the number of extra-task pairs increases, the performance of *NN-MT* increases, then tends towards that of *RN-MT*, and then degrades at larger values of 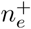 because too many dissimilar extra-task pairs are included in the training set. This implies that only a limited number of the closest extra-task pairs is required to reach optimal performance. Adding the same number of negative extra-task pairs 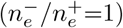 provides the best AUPR, which again limits the size of the required training set. Unsurprisingly, the best AUPR in the absence of the closest intra-task pairs (around 0.87 for *θ* = 0.80) is still lower than when all available intra-task pairs are used (AUPR=0.93, see Section 3.4). Note that, although the performance of *MT-intra* can be biased when considering similarity thresholds of 20th and 80th percentile, because the corresponding sizes of the train sets might be different, this is not the case for the *NN-MT* method because the prediction is driven by extra-task pairs.

On the contrary, Fig 6(c) and (d) show that the performance does not improve when the extra-task training pairs are chosen at random, and therefore, are on average further from the test pair. It might even degrade when the number of extra-task pairs becomes large. Finally, Fig 7 shows that, for all similarity thresholds, the performance of the *MT-intra* and *RN-MT* methods are similar, and far beyond that of the *NN-MT* method.

In conclusion, in settings where (protein, ligand) pairs similar to the test pair are available, our results suggest the best prediction performance are obtained using the *NN-MT* method trained with 10 times more negative intra-task pairs than positive ones, 1 to 5 extra-task nearest neighbor positive pairs, and the same number of extra-task negative pairs. The computational time will be reasonably comparable to that of *ligand-based ST*, and performance should be high enough (AUPR above 0.85) to guide experimental evaluations for drug specificity prediction.

**Fig 6.** AUPR score of *NN-MT* **and** *RN-MT* **as a function of the** 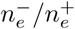 **ratio**, for a number of extra-task positive pairs 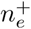 varying from 0 to 50, and for percentile-based similarity threshold *θ* of 20 and 80 applied to the intra-task positive pairs. (a): *NN-MT, θ* = 0.20. (b): *NN-MT, θ* = 0.80. (c): *RN-MT, θ* = 0.20. (d): *RN-MT, θ* = 0.80. Numerical values can be found in Supporting Information, respectively S9-S12 Tables

**Fig 7.** AUPR score as a function of percentile-based similarity *θ*, for 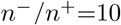, a number of extra-task positive pairs 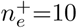 and a ratio of 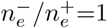 for extra-task pairs. Numerical values can be found in Supporting Information S13 Table

### 3.5.3 Adding dissimilar extra-task positive examples

While we previously argued that the point of multi-task approaches is to leverage similar extra-task data to improve prediction performance, ligand specificity studies can require the prediction of interactions between proteins and ligands for which very little similar extra-task data is available. We therefore repeated the experiments from the previous section, but this time applying the percentile-based similarity constraint to both intra-task and extra-task positive pairs of the train set. We report the corresponding *LOO-CV* AUPR on Fig 8.

We observe that the performance of *NN-MT* remains overall low (best AUPR score of 0.75 for *θ* = 0.80). Adding dissimilar extra-task positive pairs fails to improve the scores obtained when only intra-task positive pairs are included in the training set. Hence, if neither close intra-task nor close extra-task positive pairs are available, no method can provide performance good enough for the purpose of drug sensitivity prediction. These interactions would have to be experimentally tested if they are critical in the context of a drug’s development program. These observations were expected given that adding random extra-task training pairs, possibly far from the tested pair, did not improve performance of the *MT-intra* method (see Section 3.5.2).

**Fig 8.** AUPR scores of the *NN-MT* **and** *RN-MT* **multi-task methods as a function of the 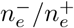 ratio**, for a number of extra-task positive pairs 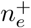 varying from 1 to 50. (a): *NN-MT, θ* = 0.20. (b): *NN-MT, θ* = 0.80. (c): *RN-MT, θ* = 0.20. (d): *RN-MT, θ* = 0.80. The two methods are trained with intra-task and extra-task examples that are both dissimilar to the tested pair (percentile-based similarity thresholds *θ* of 20 and 80). Exact values can be found in Supporting Information respectively in S14-S17 Tables

Taken together, our results show that the proposed *NN-MT* method is the most appropriate for predicting the specificity of a molecule. Indeed, it outperforms all its comparison partners independently of the number of known (protein, ligand) interacting pairs involving the same or similar ligands or proteins as the query pair. In addition, it requires much fewer training pairs than the classical *MT* approach, and its computational time is therefore close to that of a single-task method. Finally, in the most challenging setting where no similar intra-task nor extra-task training data is available, it performs significantly better than random, in a context where *ligand-based ST* could not make any prediction.

The results we have presented so far address the issue of using kerrnel methods with SVM in the context of proteome-wide specificity prediction, at a tractable computational cost thanks to the choice of a reduced learning dataset, without loss in prediction performance.

However, another key issue corresponds to study the specificity of a molecule within a family of related proteins. Indeed, when a new drug candidate is identified against a given therapeutic target, proteins belonging to the same family are important off-target candidates. This corresponds to the setting where similar training pairs are available, since proteins of the same family are similar in terms of sequence.

In the next section, we therefore assess whether the proposed *NN-MT* method, initially dedicated and tuned in proteome-wide prediction problems, also provides good performance for molecule specificity prediction within a family of proteins.

### 3.6 Specificity prediction within families of proteins

We considered three families of proteins because they gather a wide range of therapeutic targets, and have also been considered in other chemogenomics studies, thus providing reference prediction scores: G-Protein Coupled Receptors (GPCRs), ion channels (IC), and kinases. All the (protein, molecule) pairs involving GPCRs, ICs, or kinases that were present in the dataset *S* described in Section 2.4 were used to build the three corresponding family datasets.

We compared the performance of the *MT-intra* method (trained using only positive pairs involving the protein or the ligand of the tested pair) to those of the *NN-MT* and *RN-MT* methods, in order to evaluate the interest of the multi-task approach in family studies. We considered two versions: one in which the Profile protein kernel is used, as in the above sections, and another in which a family-based hierarchy kernel is used (Section 2.1), because a sequence-based kernel may not be optimal to study the specificity of the molecule within a family of proteins [11, 27]. The corresponding methods are called *MT-intra-family, NN-MT-family*, and *RN-MT-family*.

As in the above section, each (protein, ligand) test pair is considered in turn in a *LOO-CV* scheme. We used a learning dataset containing: all positive intra-task positive pairs, ten times more random negative intra-task pairs (this value was found adequate in previous sections), a varying number of positive extra-task pairs (nearest neighbors for *NN-MT* or *NN-MT-family*, random for *RN-MT* or *RN-MT-family*), and a number of negative extra-task pairs so that the ratio of 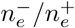 varies from 1 to 20.

#### 3.6.1 G-Protein Coupled Receptor family

Fig 9 shows that all methods perform very well, with AUPR scores above 0.95. Including extra-task positive pairs in the train set improves the AUPR score, even when added randomly. This indicates that, contrary to studies in larger scales in the protein space, in family studies, extra-task pairs are always close to the tested pair because they belong to the same family. However, the performance reached when adding positive nearest-neighbor extra-task pairs remains above those reached when adding positive random extra-task pairs, as observed in the larger scale studies presented above.

Overall, adding 10 to 50 extra-task positive pairs to the train set, and around 10 times more random negative extra-task pairs leads to the best performance.

The best AUPR scores of the *NN-MT* and the *NN-MT-family* methods are close (0.96 and 0.97). Although the best scores of the *NN-MT-family* method are slightly above those of *NN-MT*, one should note that the family GPCR kernel is based on a GPCR hierarchy that was established using known GPCR ligands. Therefore, the results obtained by the *NN-MT-family* might be biased, which is not the case for those obtained by the *NN-MT*.

**Fig 9.** AUPR score of the considered multi-task methods on the GPCR family as a function of the 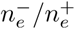 **ratio**, for a varying number 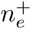 of extra-task positive pairs. (a): *NN-MT-family* (family hierarchy kernel). (b): *NN-MT* (sequence kernel). (c): *RN-MT-family* (family hierarchy kernel). (d): *RN-MT* (sequence kernel). The blue horizontal line corresponds to the *MT-intra* method trained only on intra-task pairs. Numerical values can be found in Supporting Information, respectively S18-21 Tables

#### 3.6.2 Ion Channel family

The conclusions obtained above in the GPCR family also hold in the IC family, as shown in Fig 10. Again, all methods perform very well, reaching AUPR scores above 0.97. As for the GPCR family, adding 10 to 50 extra-task positive pairs to the train set, and around 10 times more random negative extra-task pairs leads to the best performance.

**Fig 10.** AUPR of the multi-task methods on the IC family. (a): *NN-MT-family* (family hierarchy kernel). (b): *NN-MT* (sequence kernel). (c): *RN-MT-family* (family hierarchy kernel). (d): *RN-MT* (sequence kernel). The blue horizontal line corresponds to the *MT-intra* method trained only on intra-task pairs. Numerical values can be found in Supporting Information, respectively S22-25 Tables

#### 3.6.3 Kinase family

In the kinase family, the results are somewhat different from those obtained on IC and GPCRs. The *NN-MT* and *RN-MT* methods both outperform the *MT-intra* method that is trained using only intra-task pairs, as shown in Figs 11(b) and (d). Again, 10 to 50 extra-task pairs, with a 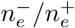 ratio in the range of 1 to 5 leads to the best results, with AUPR scores in the range of 0.93. Unexpectedly, the *NN-MT-family* and *RN-MT-family* methods, which both use the kinase family hierarchy kernel, tend to perform not as well when extra-task pairs are added to the training set, than when only the intra-task pairs are used, as shown in Figs 11(a) and (c). In addition, their best AUPR scores reaches 0.90, which is lower than those of the *NN-MT* and *RN-MT* methods which are in the range of 0.93. These observations may reflect the fact that the kinase family gathers proteins that are relatively more diverse than GPCRs and IC. For example, one can distinguish Tyrosine kinases and Serine/Threonine kinases, or globular protein kinases and receptor protein kinases. This diversity is illustrated by the organization of the kinome in some 50 distinct sub-families [43]. In this context, the sequence kernel that was optimized in proteome-wide studies might better capture the degree of similarity between two kinases than the hierarchy kernel does.

Overall, the above results on the IC, GPCR and kinase families indicate that the proposed *NN-MT* method leads to the best results when the train set includes all positive intra-task pairs, 10 times more random negative intra-task pairs, a small number of nearest neighbors positive extra-task pairs (in the range of 10) and around 5 times more random negative extra-task pairs. These conditions are very similar to those leading to the best prediction scores when ligand specificity is studied on large scale in the protein space. Even if the performance of *NN-MT* on family datasets is better than those reached by other methods on similar datasets [11, 20], they remain in the same order of magnitude.

**Fig 11.** AUPR score of the multi-task methods within the kinase family. (a): *NN-MT-family* (family hierarchy kernel). (b): *NN-MT* (sequence kernel). (c): *RN-MT-family* (family hierarchy kernel). (d): *RN-MT* (sequence kernel). The blue horizontal line corresponds to the *MT-intra* method trained only on intra-task pairs. Numerical values can be found in Supporting Information, respectively S26-S29 Tables

## 4 Discussion: comparison to other methods

As mentioned in Section 1.2, a few methods have been proposed to predict interactions between proteins and ligands. We compared the prediction performances of the proposed *NN-MT* method to those of two state-of-the art methods: a recent Matrix Factorization method called Neighborhood Regularized Logistic Matrix Factorization (*NRLMF*) [24], and the Kronecker (kernel) Regularized Least Square regression method *KronRLS* (a kernel-based method, as *NN-MT*) [18, 19].

The *KronRLS* and *NRLMF* methods were published based on their prediction performance on four protein family datasets, Nuclear Receptors (NR), GPCR, Ion channels (IC), and Enzymes (E) that contained respectively 90, 636, 1476 and 2926 interactions [10].

The *KronRLS* method uses a kernel *K*_*molecule*_ for molecules that is defined by:

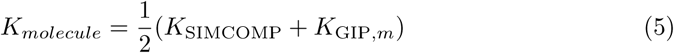

where *K*_SIMCOMP_ is a structure similarity kernel [61], and where *K*_GIP, *m*_ is a Gaussian kernel that compares the interaction profiles of molecules against the proteins of the dataset [18]. For proteins, the kernel *K*_*protein*_ is defined by:

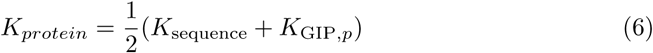

where *K*_sequence_ is a protein sequence similarity kernel also based on the Smith-Waterman score [41], and *K*_GIP,*p*_ is a Gaussian kernel that compares the interaction profile of proteins against the molecules of the dataset.

A specific feature of the *NRLMF* method is that it integrates a neighborhood regularized method which allows to take into account only the K nearest neighbors to predict a given (protein, ligand) interaction (in practice, the authors used K=5).

We performed benchmark experiments on these family datasets using the PyDTI package. This package initially contained the *KronRLS*, a variant of it called *KronRLS-WNN*, and the *NRLMF* methods, and the kernel matrices *K*_*protein*_ and *K*_*molecule*_ calculated for the four family datasets. In all experiments, we compare the intrinsic performances of the algorithms: the similarity measures used are the same for all methods. More precisely, the three methods used the kernels available in PyDTI: the structure-based *K*_SIMCOMP_ for molecules, and a kernel *K*_sequence_ based on the Smith-Waterman score. In addition, *KronRLS* also used the *K*_GIP,*m*_ kernel, leading to the *K*_*molecule*_ kernel for molecules defined in eq. 5, and the *K*_GIP,*p*_ kernel, and to the *K*_*protein*_ kernel for proteins defined in eq. 6. The two other methods do not use the *K*_GIP_ kernels because they do not take into account information about interaction profiles.

We also performed benchmark experiments on a dataset gathering more diverse protein and ligands. To this end, we used the DrugBank-based dataset *S*_0_ described in Section 2.4) containing 5 908 interactions. Because *KronRLS* and *NRLMF* could not make predictions on *S*_0_ at a manageable computational cost in the *LOO-CV* scheme, we randomly sampled 2 000 of the 5 908 interactions of this data set to create a smaller test data set called *S*_0,2000_. We still used all of *S*_0_ (minus the test example) for training.

We calculated the Tanimoto and Profile kernels optimized in the present study (see Section 3.1), and these matrices were uploaded in PyDTI so that the three considered methods could used them. In addition, since *KronRLS* also use the *K*_GIP,*m*_ and the *K*_GIP,*p*_ kernels, we described all molecules and proteins in *S*_0_ by their interaction profile. We calculated the *K*_GIP,*m*_ and *K*_GIP,*p*_ kernels on *S*_0_ and uploaded these kernels in PyDTI. Only *KronRLS* used these additional kernels. All cross-validation experiments were performed building test sets from *S*_0,2000_ and using all remaining data points in *S*_0_ for training.

Table 3 presents the performance of the three considered methods on the protein family datasets.

**Table 3.** AUPR scores and standard deviations in *10-fold-CV*, test sets balanced in positive and randomly chosen negative samples

| Method / Dataset | NR | GPCR | IC | E |
| --- | --- | --- | --- | --- |
| *KronRLS* | $0.75 \pm 0.14$ | $0.9 \pm 0.03$ | $0.96 \pm 0.01$ | $0.96 \pm 0.01$ |
| *NN-MT* | $0.89 \pm 0.09$ | $0.95 \pm 0.02$ | $0.97 \pm 0.01$ | $0.97 \pm 0.01$ |
| *NRLMF* | $0.96 \pm 0.04$ | $0.96 \pm 0.02$ | $0.98 \pm 0.01$ | $0.98 \pm 0.0$ |

Globally, the performance of all methods are high and close, with AUPR scores above 0.9 in most of the cases. On average, the *NRLMF* and *NN-MT* methods are on par and lead to the best results. These results are consistent with those reported in [24].

Table 4 confirms the tendencies observed in Table 3. Although the performances are slightly lower on this more diverse *S*_0,2000_ dataset than on the family datasets, they remain high, with NRMLF and *NN-MT* keeping the best AUPR scores.

**Table 4.**
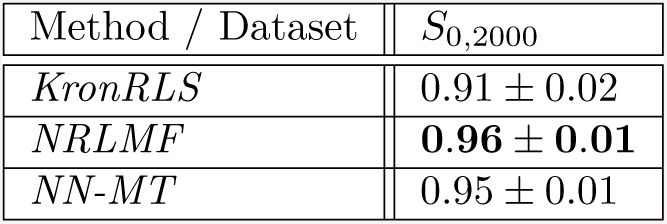
AUPR scores and standard deviations in *10-fold-CV*, test sets balanced in positive and randomly chosen negative samples

As discussed in Sections 3.1 and 3.2, various prediction methods lead to such high performances because, in the protein family or *S*_0,2000_ datasets, predictions are averaged over test pairs in which the protein and/or the molecule might be orphan, or not. These averaged results hide less favorable situations, typically double orphan samples. Because these cases are common and important when predicting specificity of a new drug candidate at the proteome scale, we would like to stress that comparing methods in orphan cases is a more stringent and relevant test. In such cases, the performance are expected to be more modest and the methods might not rank in the same order.

Therefore, we ran the three methods using a *LOO-CV* scheme on double orphan (protein, molecule) pairs on the same datasets. In these experiments, for each tested (*p, m*) pair, interactions involving the considered protein or the molecule are ignored in the train set. The *LOO-CV* schemes were balanced in positive and randomly chosen negative pairs.

In order to better explore the performance of the considered methods in this double-orphan setting, we used two versions of the two kernel-based methods, initially introduced for *KronRLS*. More precisely, in [19], the authors proposed an approach called WNN (weighted nearest neighbor) that, for each orphan molecule *m* (resp. protein), an interaction profile is computed by summing the weighted profiles of non orphan molecules in the dataset. The weighting depends on the similarity between the orphan molecule and all other non orphan molecules. This predicted profile is used in the training to predict labels to all (protein, *m*) pairs of the dataset. Thus, in the first version of *KronRLS* [18], *all the labels of (protein, m*) pairs involving the orphan molecule *m* were set to 0. Based on this WNN procedure some of these non interactions might be requalified as true interaction before training the predictor. In other words, the WNN algorithm can be viewed as a mean to de-orphanize molecules or proteins in order to help the predictions on such cases. In the following, we will call *KronRLS-WNN* the *KronRLS* method ran with the WNN algorithm. Using the PyDTI package, we also considered a version of *NN-MT* in which the WNN algorithm is the implemented, and call it *NN-MT-WNN* in the following.

Table 5 presents the results of the double-orphan benchmark on the family datasets. Surprisingly, in these double-orphan experiments, the NRMLF method has very modest results and does not perform as well as the other methods. The results of *NN-MT* remain well above the random performance of 0.5, but not the *KronRLS* method. The WNN algorithm dramatically improves the performance of *KronRLS*, and to a lesser extent those of *NN-MT*, and overall, the *NN-MT-WNN* algorithm leads to the best performance in most cases.

**Table 5.** AUPR scores and standard deviations on double orphan *LOO-CV*, balanced number of positive and randomly chosen negative test samples

| Method / Dataset | NR | GPCR | IC | E |
| --- | --- | --- | --- | --- |
| *KronRLS-WNN* | $0.78 \pm 0.03$ | $0.83 \pm 0.01$ | $0.78 \pm 0.01$ | $0.84 \pm 0.0$ |
| *KronRLS* | $0.55 \pm 0.01$ | $0.62 \pm 0.01$ | $0.64 \pm 0.0$ | $0.56 \pm 0.0$ |
| *NRLMF* | $0.19 \pm 0.03$ | $0.15 \pm 0.0$ | $0.26 \pm 0.01$ | $0.23 \pm 0.0$ |
| *NN-MT* | $0.72 \pm 0.04$ | $0.76 \pm 0.01$ | $0.72 \pm 0.01$ | $0.66 \pm 0.0$ |
| *NN-MT-WNN* | **$0.77 \pm 0.05$** | $0.85 \pm 0.0$ | $0.79 \pm 0.01$ | $0.84 \pm 0.0$ |

Table 6 presents the results of the double-orphan benchmark *S*_0,2000_ dataset. We did not run the *NRLMF* method in this experiment, because it was computationally too intensive in this *LOO-CV*, and because it already gave very poor results on the easier family dataset. Moreover we shortened the train set of *KronRLS* and *KronRLS-WNN* methods by considering only the thousand molecules (resp. proteins) closest to the molecule (resp. protein) of the test sample. Thus, the computation time was reduced to some hours instead of months which made those methods computationally reasonable.

**Table 6.** AUPR scores and standard deviations on double orphan *LOO-CV*, balanced number of positive and randomly chosen negative test samples

| Method / Dataset | $S_{0,2000}$ |
| --- | --- |
| *NRLMF* | None |
| *KronRLS-WNN* | $75.6 \pm 0.43$ |
| *KronRLS* | $0.4979 \pm 0.0071$ |
| *NN-MT* | $0.60 \pm 0.01$ |
| *NN-MT-WNN* | **$0.85 \pm 0.01$** |

Overall, the scores are lower on this dataset than on the family datasets because *S*_0,2000_ is a more diverse dataset on which predictions are more difficult, in general, than on the family datasets. However, the same tendency is observed: *NN-MT* performs better than *KronRLS*, and when the WNN algorithm is used, *NN-MT-WNN* performs better than *KronRLS-WNN*.

Overall, the results of these benchmarks show that the *NN-MT* method present state-of-the-art or better results on the protein family datasets and the more diverse DrugBank-based dataset. In the general case, it appears to be a good default method in terms of performance, number of parameters and computational efficiency, which are important issues for non expert users.

In the specific double-orphan case, only the two kernel-based methods *NN-MT* and *KronRLS* lead to performance well above those of a random predictor. The WNN algorithm, proposed in [19] improves the performance of *KronRLS* and of *NN-MT*, but resulting *NN-MT-WNN* method lead to the best performance.

Finally, it is interesting to compare the computational complexities of the methods as a function of the number of hyper-parameters that they contain. Indeed, these hyper-parameters need to be optimized by cross-validation, leading to heavy computational issues in the case of the large-scale datasets used in proteome-wide chemogenomics. As can be seen in the PyDTI package, *NRLMF* has 5 regularization parameters, *KronRLS* has 2 hyper-parameters (decay parameter T and the weight parameter used to combine kernels; the regularization parameter and the bandwidth of the GIP kernel are fixed), and *NN-MT* has 1 hyper-parameter (regularization parameter *C* for SVM). In practice, the optimization of *NRLMF* in the *LOO-CV* scheme was out of reach, requiring several days of calculation while the other methods required hours. This could explain in part the very low performances displayed by *NRLMF* in the double-orphan experiment. However, we did cross-validate *NRLMF* parameters in the double-orphan setting in the case of the family NR dataset (the smallest dataset used in this section). This allowed a modest increase in AUPR score from 0.14 to 0.19 (reported in Table 5). Therefore, even if the *NRLMF* method had been optimized on the other datasets, we do not expect that this would have changed the overall conclusion that this method is not suitable for handling orphan cases.

## 5 Conclusion

The present study tackles prediction of ligand specificity on large scale in the space of proteins. More precisely, our goal was to propose a method to explore the specificity molecules with state-of-the-art or better performance over a wide range of prediction situations: at the proteome or protein family scales, on average or in specific situations such as tested pairs far from the train set, or such as orphan proteins and ligands. In other words, the aim was to propose a robust default method, applicable to many types of studies, thus avoiding development of *ad hoc* complex and specific methods to non expert users. We chose to formulated it as a problem of predicting (protein, ligand) interactions within a multi-task framework based on SVM and Kronecker products of kernels on proteins and molecules. Within the kernel-based SVM methods tested in the Results section, we showed that the *NN-MT* method fulfills these requirements. In particular, *NN-MT* outperforms both the multi-task *MT* method and the corresponding single-task kernel-based methods, while it also keeps a computational cost close to that of single-task approaches. The *NN-MT* algorithm fulfills these requirements, leading to the best prediction performance for the three tested settings which cover most of the prediction situations that would be encountered in real-case studies.

To summarize the main characteristics of the proposed *NN-MT* method (detailed in Sections 3.1, 3.4 and 3.5), we suggest to predict each (protein, ligand) interaction using the Profile kernel for proteins (with subsequences length k of 5 and threshold equaled to 7.5) and the Tanimoto kernel for molecules (with length of path 8), with a train set including:

- all positive intra-task pairs (i.e. all known interactions involving the protein or the ligand of the test pair), and around ten times more randomly chosen intra-task negative pairs.
- a small number of the closest positive extra-task pairs (i.e. a number similar to that of intra-task positive pairs), and a similar number of randomly chosen negatives extra-tasks pairs.

This should provide good default parameters to use the *NN-MT* method in a straightforward manner for users that are not familiar with machine learning approaches.

Overall, the *NN-MT* method could be an interesting tool to guide not only the choice of the best hit molecules early in the drug development process avoiding deleterious side effects for patients, but also to suggest drug repositioning opportunities since a side effect for a patient might be viewed as a therapeutic effect for another.

We used the DrugBank database to build several datasets that illustrate various prediction contexts and that we made available online to the community for future benchmarking studies. We also updated the PyDTI package [24] with an implementation of *NN-MT* together with several cross-validation schemes and our DrugBank-based dataset. This allowed us to compare the *NN-MT* method to recent approaches developed on drug target interaction prediction [18–24]. In the context of wide-scale prediction of molecule specificity, the DrugBank-based dataset is more relevant that the family datasets that have been widely used. Indeed, it contains a set of proteins that can be viewed as a relevant druggable proteome to train and test computational models for drug specificity prediction.

The benchmark study comparing *NN-MT* to the matrix factorization NRMLF and the kernel-based *KronRLS* algorithms on family and DrugBank-based datasets showed that, all methods displayed high performances, NRMLF and *NN-MT* leading to the best results. However, on the more demanding double-orphan tests performed on the same datasets, NRMLF performed much poorer than the kernel-based *NN-MT* and *KronRLS* algorithms. In this orphan case, the WNN algorithm makes it possible not only to significantly improve the performance of *KronRLS*, but also that of *NN-MT*, the *NN-MT-WNN* algorithm leading to the best results.

We formalized (protein, ligand) interaction prediction as a classification problem because they can be solved at a reasonable computational cost on large datasets. Whenever the purpose would be to predict the relative affinities of molecules for a set of proteins, the predicted scores can be used to rank all interactions. However, this question could be also solved using a regression algorithm when predicting the affinity between pair of molecule and protein [26]. Note that in such an approach, the affinity of all (protein, ligand) pairs in the training data is required, which is rarely available on large scale.

Although the protein-ligand interaction process takes place in the 3D space, we chose to encode the two partners based on features that do not require 3D information. Indeed, the bound conformation of the ligand, and the 3D structure of the protein binding pocket is unknown in most cases, which prevents predictions on large scale. However, we are aware that a method using 3D information to encode the interaction can be of interest on more restricted datasets (i.e. not covering the druggable human proteome) as those available from the PDB database [28, 29].

Since the prediction performance strongly depends on the distance between the predicted interactions and the train set, it could be relevant to apply the multi-kernel learning (MKL) framework [62]. Indeed, different feature spaces will lead to different metrics which could modulate the distance between the test and train sets. This idea was explored in [63], but in this work the MKL approach employed L2 regularization between kernels, which did not lead to the improvements that could be expected from an L1 (i.e. sparsity-inducing) regularization term.

For future developments of the method, it is likewise relevant to explore the benefits of deep learning approaches in the context of representation learning [64]. Indeed, learning the featurization of molecules [65] and of proteins [66] on various prediction tasks including drug-target interaction prediction could optimize the featurization for the specificity prediction task. The present work showed that learning on structurally similar compounds and similar proteins (according to sequence similarity) improves and speeds up the prediction performance on drug-target interaction task. A recent study [67] showed that in the case of stacked fully connected layers, learning with structurally similar compounds but uncorrelated activities can provide contradictory information leading to a decrease of performance. Even more recently, it was shown that current graph-CNN based models perform best when trained on compounds similar to the tested compounds [68], as we observed for the *NN-MT* method proposed in the present study. However, deep learning based models seem promising to efficiently share information between more dissimilar compounds and putative targets as they can actually learn a generic representation of molecules and proteins based on several supervised learning prediction task.

## Supporting information

**S1 Fig. (A) Scores of** *MT* **Kernel Ridge regression on datasets S1/2/3 with a** *nested 5-fold-CV* **scheme. (B) Scores of** *MT* **Kernel Ridge regression on S1 depending on the CV scheme.**

**S2 Fig. Scores of the** *MT* **method on S1 depending on CV scheme.** Overall, all CV schemes provide high prediction performance on this dataset, in the range of 0.93-0.94 in AUC and AUPR. The *nested 5-fold-CV* leads to performance very close to those of *5-fold-CV*, showing that on the S1 dataset, *5-fold-CV* did not suffer from overestimation of the performance due to data over-fitting. *LOO-CV* leads to slightly better results, although very close to those of the other CV schemes. In general, the *LOO-CV* scheme is expected to provide better results because the model is trained on more data points than in *5-fold-CV*. Again, this problem seems to be limited here, since the performance of *LOO-CV* does not differ much from that of *nested 5-fold-CV*.

**S1 Table. scores of** *MT* **SVM in** *nested 5-fold-CV* **scheme on S1–S4 datasets.**

**S2 Table. scores of** *MT* **SVM with** *nested 5-fold-CV* **scheme on S1’–S4’ datasets.**

**S3 Table. scores of** *MT* **SVM on S1 dataset, depending on the CV scheme.**

**S4 Table. scores of** *ligand-based ST* **and** *MT-intra* **methods with the** *LOO-CV* **scheme on S1 dataset.**

**S5 Table. scores of** *NN-MT* **in** *LOO-CV* **scheme on S1 dataset.**

**S6 Table. scores of** *RN-MT* **in** *LOO-CV* **scheme on S1 dataset.**

**S7 Table. scores of** *MT-intra* **in** *LOO-CV* **on S1 dataset with similarity constraint on intra-task pairs.**

**S8 Table. scores of** *ligand-based ST* **in** *LOO-CV* **scheme on S1 with similarity constraint on intra-task pairs.**

**S9 Table. scores of** *NN-MT* **in** *LOO-CV* **scheme on S1 dataset, with similarity constraint on intra-task pairs (***θ* = 20).

**S10 Table. scores of** *NN-MT* **in** *LOO-CV* **scheme on S1 dataset, with similarity constraint on intra-task pairs (***θ* = 80).

**S11 Table. scores of** *RN-MT* **in** *LOO-CV* **on S1 with similarity constraint on intra-task pairs (***θ* = 20).

**S12 Table. scores of** *RN-MT* **in** *LOO-CV* **on S1 with similarity constraint on intra-task pairs (***θ* = 80).

**S13 Table. scores of** *MT-intra, NN-MT, RN-MT* **in** *LOO-CV* **on S1 dataset, with similarity constraint on intra-task pairs.**

**S14 Table. scores of** *NN-MT* **in** *LOO-CV* **on S1, with similarity constraint on intra- and extra-task pairs (***θ* = 20)

**S15 Table. scores of** *NN-MT* **in** *LOO-CV* **on S1, with similarity constraint on intra- and extra-task pairs (***θ* = 80)

**S16 Table. scores of** *RN-MT* **in** *LOO-CV* **on S1 dataset, with similarity constraint on intra- and extra-task pairs (***θ* = 20)

**S17 Table. scores of** *RN-MT* **in** *LOO-CV* **on S1 dataset, with similarity constraint on intra- and extra-task pairs (***θ* = 80)

**S18 Table. GPCR dataset: scores of** *NN-MT* **in** *LOO-CV* **scheme with family’s hierarchy based kernel**

**S19 Table. GPCR dataset: scores of** *NN-MT* **in** *LOO-CV* **scheme with sequence based kernel**

**S20 Table. GPCR dataset: scores of** *RN-MT* **in** *LOO-CV* **scheme with family’s hierarchy based kernel**

**S21 Table. GPCR dataset: scores of** *RN-MT* **in** *LOO-CV* **scheme with sequence based kernel**

**S22 Table. Ion Channel dataset: scores of** *NN-MT* **in** *LOO-CV* **scheme with family’s hierarchy based kernel**

**S23 Table. Ion Channel dataset: scores of** *NN-MT* **in** *LOO-CV* **scheme with sequence based kernel**

**S24 Table. Ion Channel dataset: scores of** *RN-MT* **in** *LOO-CV* **scheme with family’s hierarchy based kernel**

**S25 Table. Ion Channel dataset: scores of** *RN-MT* **in** *LOO-CV* **scheme with sequence based kernel**

**S26 Table. Kinase dataset: scores of** *NN-MT* **in** *LOO-CV* **scheme with family’s hierarchy based kernel**

**S27 Table. Kinase dataset: scores of** *NN-MT* **in** *LOO-CV* **scheme with sequence based kernel**

**S28 Table. Kinase dataset: scores of** *NN-MT* **in** *LOO-CV* **scheme with family’s hierarchy based kernel**

**S29 Table. Kinase dataset: scores of** *NN-MT* **in** *LOO-CV* **scheme with sequence based kernel**

## S1 Appendix. Basic principles of SVM

Let us consider a set of labeled samples S = *{*(*x*_1_, *y*_1_), *…,* (*x*_*N*_, *y*_*N*_)*}* where (*x*_*i*_, *y*_*i*_) *∈ X ×* {-1, +1} for *i* = 1, *…, N*, and where the space *X* into which the data points live is equipped with a dot product 〈…, 〉 For example, the data points *x*_*i*_ represent ligands, and their labels *y*_*i*_ are equal to +1 for ligands that bind to a given protein and −1 for ligands that don’t. In the simplest case where the two classes of data points are linearly separable, Support Vector Machines [32] (SVM) is an algorithm that learns to separate these two classes based on an hyperplane whose equation can be defined by a normal vector w and a constant b: 〈*w, x〉* + *b* = 0. Among the infinity of potential separating hyperplanes, the optimal hyperplane maximizes the margin. This margin is defined as the closest distance from the hyperplane to any of the data points. It can be shown that the search of this optimal hyperplane can be formulated by the following optimization problem:

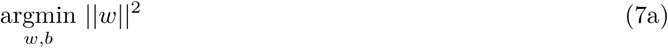

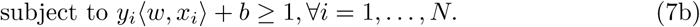

The solution hyperplane maximizes its distance to the closest data points, and this distance is equal to 2*/‖w‖*^2^.

Then, the decision function f allowing to make predictions for any new point x depends on its position with respect to the hyperplane, i.e. based on the sign of *f* (*x*) = 〈*w, x*〉 + *b*.

This optimization problem is strictly convex and admits a unique solution. The Lagrangian associated to the optimization problem leads to the following equivalent dual problem:

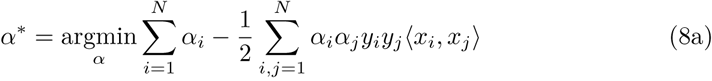

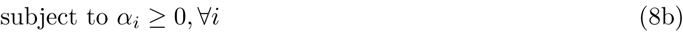

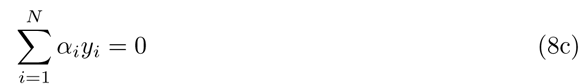

where the coefficients *α*_*i*_ are known as the Lagrange multipliers associated to the constraints *y*_*i*_ 〈*w, x*_*i*_ 〉 + *b ≥* 1.

In practice, this quadratic problem that can be solved efficiently using a dedicated algorithm, known as Sequential Minimal Optimization (SMO) [69]. When the optimum *α*^*\**^ is met, the decision function allowing to make predictions for any new point x depends on its position with respect to the hyperplane:

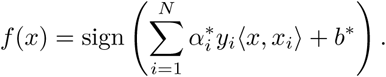

However, the two classes of data points may not be linearly separable. In these situations, kernel methods are a widely-used set of techniques that allow to adapt linear methods to non-linear models. Let us consider a semi-definite positive kernel function *K*: *X × X → R*. The Mercer theorem states that there exists a non-linear function *ϕ*: *X → H* that maps data points in *X* into a high dimensional feature Hilbert space *H* where *K* can be expressed as a scalar product: *k*(*x*_1_, *x*_2_) = 〈 *ϕ*(*x*_1_), *ϕ*(*x*_2_) 〉 _*H*_. In practice, *H* is more often taken to be *R*^*d*^. Although the two classes of data points might not be linearly separable in *X*, they might become linearly separable in the high dimensional space *H* where the SVM can be solved. The principle of kernel trick is that, since the images of the data point *ϕ*(*x*_*i*_) are used only in scalar products, finding the *α*_*i*_ coefficients to solve the SVM can be done by replacing all occurrences of the scalar product 〈 *ϕ*(*x*_*i*_), *ϕ*(*x*_*j*_) *〉*_*H*_ by the kernel function *k*(*x*_*i*_, *x*_*j*_):

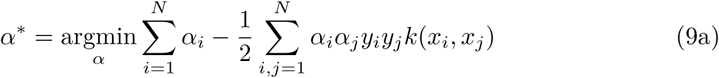

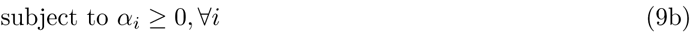

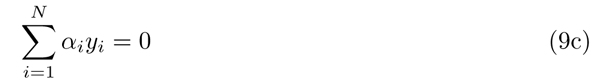

In other words, finding the separating hyperplane in *H* does not require explicit definition of the nonlinear mapping function *ϕ*, or calculation of the image vectors *ϕ*(*x*_*i*_).

Then, the label of a new data point *x* is then predicted by a function *f* (*x*) defined as:

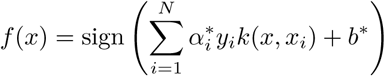

In the case where the two classes of points are not separable, we need to allow some of the training points to be misclassified, i.e. to be on the side of the separating hyperplane corresponding to points affected to the opposite label. To this end, we introduce a penalty terms *∈*_*n*_*∀n* = 1, *… N* (also called slacked variables) defined by: *∈*_*n*_ = 0 for data points that are in the correct margin boundary and *∈*_*n*_ = *|y*_*n*_ - (〈 *w, x*_*n*_〉 + *b*) *|* for the misclassified points. Thus, points on the decision boundary will have *∈*_*n*_ = 1, and misclassified points would be penalized by *∈*_*n*_ *>* 1 proportionally to their distance to the separating hyperplane. Thus, the penalty terms can be written as *∈*_*n*_ = *max*(0, 1 *y*_*n*_(〈 *w, x*_*n*_ 〉 + *b*)). Then the exact classification constraints of equation 7b are replaced by *y*_*i*_ 〈*w, x*_*i*_〉 + *b ≥* 1 *∈*_*i*_. In addition, the penalty terms must satisfy *∈*_*n*_ *≥* 0 ∀ *n* = 1, *… N*. The new objective function aims at both maximizing the margin and minimizing the penalty terms, i.e. minimizing the number of misclassified points.

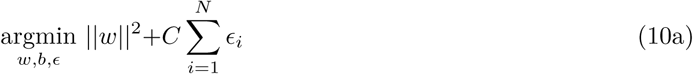

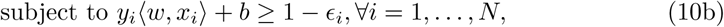

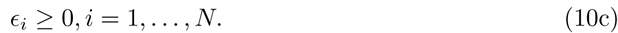

The parameter C in the objective function in equation 10a is meant to introduce a trade-off between the maximization of the margin, expressed by the term 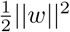, and the classification error on the training set, expressed by the penalty terms. This parameter is usually determined by cross validation on the training data. In the present study, the optimal parameter C was searched between 10^−5^ and 10^5^. As for the separable case, the SVM can also be solved in the non-separable case using a kernel function.

## S2 Appendix. Definition of the Kronecker product of two matrices A and B

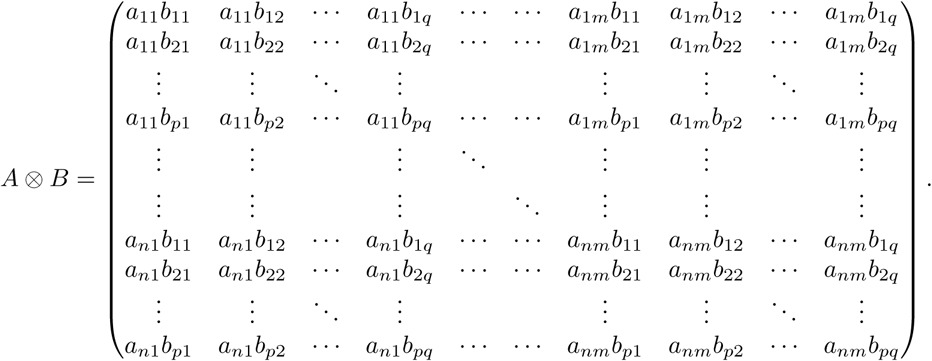

Therefore, if matrix *A* is of size *n.m* and matrix *B* is of size *p.q*, the Kronecker product of *A* and *B* is a matrix of size *n.m.p.q*

## Acknowledgments

This work was supported by the Mines ParisTech, and by Ministère de l’Industrie.

